# Development of a thermostable and broadly neutralizing pan-sarbecovirus vaccine candidate

**DOI:** 10.1101/2025.03.17.643667

**Authors:** Simran Srivastava, Sahil Kumar, Suman Mishra, Raju S. Rajmani, Randhir Singh, Somnath Dutta, Rajesh P. Ringe, Raghavan Varadarajan

## Abstract

Zoonotic spillover of sarbecoviruses to humans resulted in theSARS-CoV-1 outbreak in 2003 and the current COVID-19 pandemic caused by SARS-CoV-2. In both cases, the viral spike protein (S) is the principal target of neutralising antibodies that prevent infection. Within spike, the immunodominant receptor-binding domain (RBD) is the primary target of neutralising antibodies in COVID-19 convalescent sera and vaccine recipients. We have constructed stabilized RBD derivatives of different sarbecoviruses: SARS-CoV-1 (Clade 1a), WIV-1 (Clade 1a), RaTG13 (Clade 1b), RmYN02 (Clade 2) and BtKY72 (Clade 3). Stabilization enhanced yield by an 3-23-fold. The RBD derivatives were conformationally intact as assayed by binding to multiple broadly neutralizing antibodies. The stabilized RBDs show significant enhancement in apparent T_m_, exhibit resistance to a 2-hour incubation at temperatures up to 60℃ in PBS in contrast to corresponding WT RBDs, and show prolonged stability of over 15 days at 37℃ after lyophilization. In mice immunizations, both stabilization and trimerization significantly enhanced elicited neutralization titers by ∼100 fold. The stabilized RBD cocktail elicited high neutralizing titers against both homologous and heterologous pseudoviruses. The immunogenicity of the vaccine formulation was assessed in both naïve and SARS-CoV-2 pre-immunized mice, revealing an absence of immune imprinting, thus indicating its suitability for use in future sarbecovirus-origin epidemics or pandemics.

**Author summary:** The COVID-19 pandemic was caused by the sarbecovirus SARS-CoV-2. Phylogenetically, sarbecoviruses are divided into four clades: Clade 1a, Clade 1b, Clade 2 and Clade 3 and within these clades, are many other sarbecovirus strains with pandemic or epidemic potential. It is therefore important to develop a broadly protective, pan-sarbecovirus vaccine formulation that can be cheaply and rapidly produced. While mRNA vaccine formulations are efficacious, they have stringent low temperature storage requirements and there is limited manufacturing expertise in low and middle income countries for this modality. Neutralizing antibodies are important for protection and in the case of SARS-CoV-2 are primarily directed against the Receptor Binding Domain (RBD) of the surface spike protein. In this study, we have designed and developed an adjuvanted, protein subunit, pan sarbecovirus vaccine formulation using stabilized RBD derivatives from diverse sarbecoviruses as immunogens. We demonstrate that the stabilized RBD derivatives have considerably enhanced yield and thermal stability relative to corresponding WT proteins and that the formulation remains stable for up to two weeks at 37°C. The formulation was found to be highly immunogenic in both naïve and pre-immunized mice, eliciting neutralizing titers well above the known protective threshold, indicating its suitability for use in future sarbecovirus-origin epidemics or pandemics.

## Introduction

In both the earlier SARS-CoV-1 outbreak in 2003 and the current COVID-19 pandemic caused by SARS-CoV-2. In both cases, neutralising antibodies that prevent infection are primarily directed against the surface Spike protein (S) of the virus. Understanding and monitoring potential reservoirs, intermediary hosts, mechanisms of zoonotic transmission and focusing on pandemic preparedness are crucial for early detection and containment(1–4). In recent decades, the global community has faced unprecedented challenges posed by emerging coronaviruses, exemplified by the severe acute respiratory syndrome coronavirus (SARS-CoV) in 2002-2003, the Middle East respiratory syndrome coronavirus (MERS-CoV) in 2012, and, more recently, the ongoing coronavirus disease 2019 (COVID-19) pandemic caused by the severe acute respiratory syndrome coronavirus 2 (SARS-CoV-2)(5–11). These outbreaks have not only exposed the vulnerabilities of existing public health infrastructure but have underscored the urgent need for innovative and broadly protective vaccine strategies. The development of a broadly protective coronavirus vaccine represents an important approach to pandemic preparedness and response. Traditional vaccine efforts have often focused on specific viral strains, requiring constant adaptation to the evolving landscape of viral diversity(12,13). In contrast, a broadly protective vaccine seeks to provide immunity against a spectrum of coronaviruses, minimizing the impact of viral mutations and potential future pandemics.

During the COVID-19 pandemic, mRNA vaccines were the quickest to be deployed, but were primarily administered to those in high income countries, in part because of cost and storage temperature issues(14–17). Protein subunit vaccines offer some advantages over other vaccine platforms, contributing to their safety, efficacy, and ease of production. The receptor binding domain (RBD) of the severe acute respiratory syndrome coronavirus 2 (SARS-CoV-2) spike protein is a compelling and promising vaccine candidate and has garnered significant attention due to its pivotal role in viral entry into host cells, and its potential to elicit a robust immune response. The exploration of RBD as a vaccine target is motivated by several key factors that contribute to its potency and efficacy. The RBD is a crucial component of the spike protein responsible for binding to the angiotensin-converting enzyme 2 (ACE2) receptor on host cells(18–20). Targeting the RBD allows for interference with the initial steps of viral entry, preventing infection and replication. The RBD is known to induce a protective immune response, with a significant portion of neutralizing antibodies being directed against this domain(21–27). Vaccination with the RBD can stimulate the production of high titres of neutralizing antibodies, conferring protection against SARS-CoV-2 infection(28–33). In addition, focusing on the RBD provides a specific antigenic target, reducing the risk of immune responses against non-essential regions of the virus. Importantly, RBD exhibits a degree of amino acid sequence conservation across different SARS-CoV-2 variants and sarbecoviruses(34). This conservation suggests that vaccines targeting the RBD may confer cross-protection against pre-emergent sarbecoviruses, contributing to the long-term effectiveness of vaccination efforts. Numerous monoclonal antibodies that bind to the conserved regions of SARS-CoV-2 receptor-binding domain (RBD), such as CR3022, S309, DH1047, 10-40, S2X259 and ADG-20, have been documented to exhibit cross-neutralizing capabilities against multiple sarbecoviruses(35–40). Recent reports have shown the efficacy of multiple antigen (RBD derived from sarbecoviruses) displayed on nanoparticles to elicit broadly neutralizing response in mice as well as NHPs(41–44). In addition, multiplexed chimeric spike mRNA based vaccine also result in cross-neutralizing antibody response(45). Multivalent nanoparticle vaccine candidates have been shown to elicit protection against MERS CoV and sarbecoviruses(46–48). These studies highlight the feasibility of developing a universal sarbecovirus vaccine utilizing diverse sarbecovirus RBDs as immunogens.

We have previously demonstrated that introduction of three stabilising mutations into wild type (WT) SARS-CoV-2-RBD significantly improves thermal stability and immunogenicity(28,30). In this study, to determine the transferability of these mutation to other sarbecovirus RBDs, we introduced them in diverse RBDs from SARS-CoV-1 (Clade 1a), WIV-1 (Clade 1a), RaTG13 (Clade 1b), RmYN02 (Clade 2) and BtKY72 (Clade 3). Introduction of the mutations substantially improved thermal stability by ∼7°C, and enhanced purified yield by about 3-23 fold, relative to corresponding WT RBDs, without affecting their binding to conformation specific ligands. Cryo-EM of SARS-CoV-2 spike containing the stabilizing mutations provided some insights into the structural basis for enhanced stability. Stabilized SARS-CoV-1, WIV-1, RmYN02 and BtKY72 trimeric RBD derivatives are resistant to 2-hour incubation at temperatures of upto 60℃ in PBS, in contrast to corresponding WT RBDs. The lyophilized RBD cocktail retained antigenicity even after storage at 37°C for over 15 days and we have previously shown that lyophilized, stabilized SARS-CoV-2 RBD retains immunogenicity after month long storage at 37°C. We assessed the antigenicity of individual stabilized RBD derivatives along with the RBD cocktail formulation in mice, and comparable antibody titres were observed in both cases. In addition, significant neutralizing titers against homologous and heterologous pseudoviruses were observed in mice immunized with our stabilized sarbecovirus RBD cocktail formulation. Developing thermostable vaccines offers several significant advantages, especially in the context of low-middle income countries, where maintaining an effective cold chain during vaccine distribution can be challenging. Thermostable vaccines can be deployed quickly without the need for extensive cold chain preparations, helping to contain outbreaks and prevent the spread of infectious diseases.

## Results

### Stabilization of diverse sarbecovirus RBDs by transfer of stabilizing mutations from SARS-CoV-2 RBD

The SARS-CoV-2 RBD is part of the viral S protein, which protrudes from the virus surface and facilitates interaction with host cell receptors(18). We have previously isolated stabilized mutants of SARS-CoV-2 RBD encompassing residues 332-532 (Figure 1A). Three previously identified stabilizing mutations are A348P, Y365W and P527L (Figure 1B). Sarbecoviruses belong to the Coronaviridae family and are characterized by their large, positive single-stranded RNA genome. They have been classified into different clades based on phylogenetic analyses of their genetic sequences(49) (Figure1C). The RBD of clade 1 viruses utilize ACE2 as the receptor for cellular entry. SARS-CoV-1and SARS-CoV-2 are examples of clade-1a and clade-1b respectively, that have made the cross-species jump to infect humans(5,9,50). A specific deletion in the RBD of clade 2 viruses prevents them from using ACE2, however they do have the capacity to infect human cells through an unknown receptor(51). Interestingly, some clade 3 viruses which are found more widely in Africa and Europe can infect using ACE2 while others utilize an unknown receptor(52,53). In an attempt to select representative sarbecoviruses from each clade, we compared the % amino acid identity of the specific RBDs with SARS-CoV2 RBD (Figure 1D) and with other members of the respective clades (Figure 1E). Considering the conservation of RBD sequence within the clade as well as within SARS-CoV-2 variants, previous and ongoing outbreaks, receptor usage and potential infectivity in human cells, we selected SARS-CoV-1and WIV-1 (clade 1a); SARS-CoV-2 and RaTG13 (clade 1b); RmYN02 (clade 2) and BtKY72 (clade 3) as the representatives for their respective clades. To substantiate whether we can transfer the previously identified stabilizing mutations in SARS-CoV-2 to these diverse sarbecoviruses sharing varying degrees of amino acid identity with SARS-CoV-2 (68-90%), we analyzed the sequence and structural conservation of the target residues (Figure S1). While residue 365 and 527 were fully conserved in all the selected sarbecoviruses, it was interesting to note that one of the SARS-CoV-2 stabilizing mutations A348P was naturally present in clade 1a, 2 and 3 viruses. We therefore introduced A348P, Y365W and P527L mutations in the selected sarbecoviruses in order to study the transferability of these mutations in diverse sequence backgrounds.

**Figure 1:**
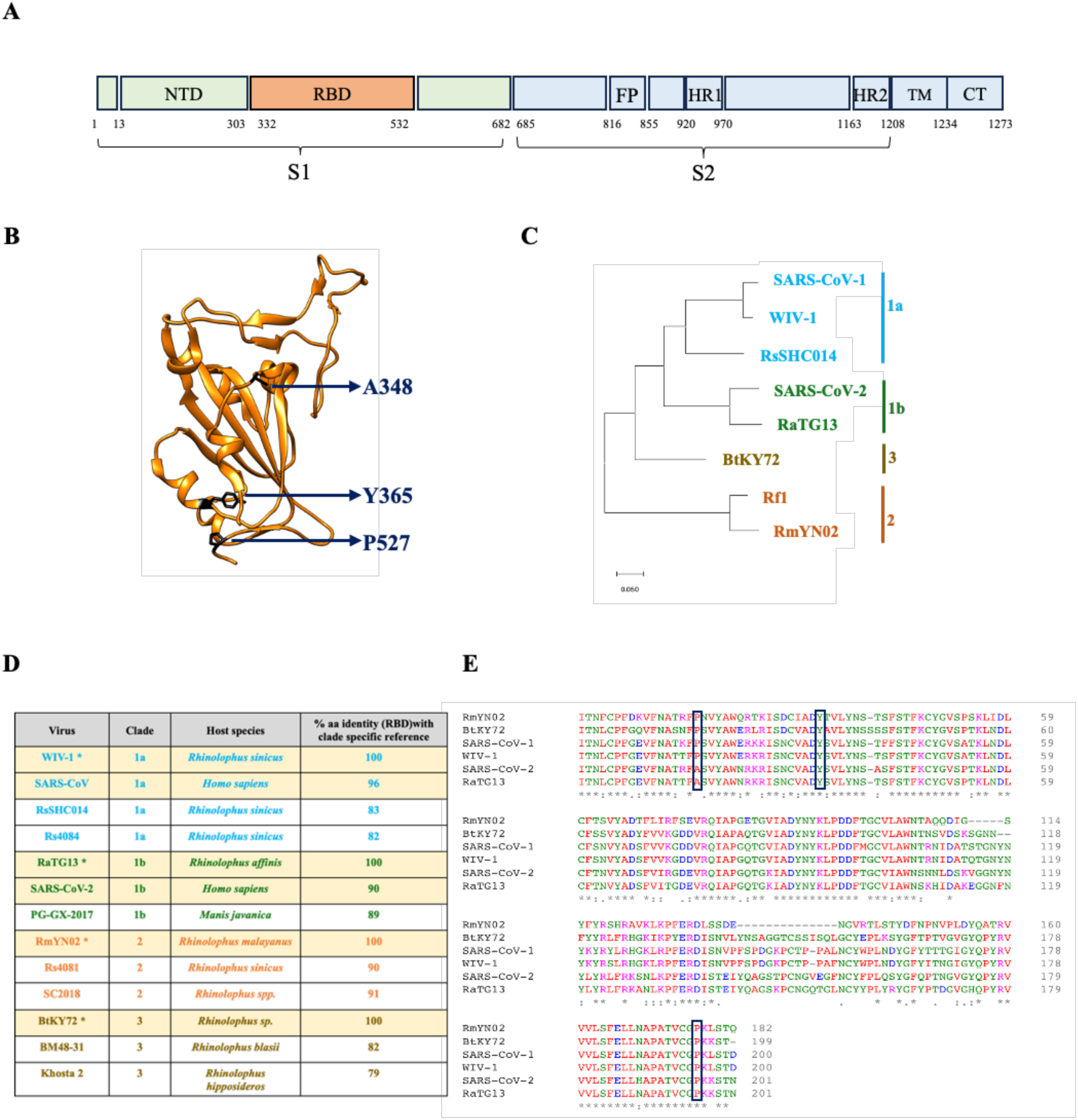
Phylogenetic analysis of sarbecovirus RBDs. (A) Schematic representation of domains in SARS CoV-2 spike protein (B) SARS CoV-2 RBD (PDB ID: 6M0J) with the residues to be mutated highlighted in the structure. (C) Phylogenetic tree of sarbecovirus RBDs constructed using MEGA software (D) % amino acid identity of selected sarbecovirus RBDs with SARS CoV-2 RBD (E) Multiple sequence alignment of sarbecovirus RBD sequences.

### Transfer of stabilizing mutations to diverse sarbecovirus RBDs results in thermal stabilization and yield enhancement without affecting conformational integrity

SARS-CoV-1 shares 74% amino acid identity with SARS-CoV-2 RBD and is closely related to another clade 1a member, WIV-1 (96% identical). WIV-1 is found in horseshoe bats (*Rhinolophus sinicus*) in China; its usage of ACE2 as the receptor highlights the potential for zoonotic transmission to humans(54). RaTG13 shares a high genetic similarity with SARS-CoV-2, it was also identified in horseshoe bats (*Rhinolophus affinis*) in Yunnan Province, China. Similar to SARS-CoV-2, RaTG13 is capable of utilizing the angiotensin-converting enzyme 2 (ACE2) receptor for cell entry.(55) RmYN02 is a recently discovered clade-2 virus which contains two deletions in the RBD that prevents it from using ACE2(56). Another SARS-related CoV that we included in this study is BtKY72 (clade-3) which was identified in Kenyan *Rhinolophus* bats and shows human ACE2-dependent entry(50,53). As oligomerization of antigen results in enhanced immunogenicity, we expressed the wild type (WT) and stabilized (St) derivatives of these sarbecovirus RBDs in *Expi-293F* cells as trimers through genetic fusion of a disulfide linked trimerization motif at the respective C termini (Figure 2A). For comparison, we also expressed and purified monomeric WT and St derivatives of SARS-CoV-1 RBD. Purification of these mammalian cell expressed proteins was performed via Ni-affinity chromatography. A significant enhancement in the yield (∼4-23-fold increase) of the RBD derivatives was observed upon introduction of the stabilizing mutations (Figure 2B). We further probed the effect of these mutations on protein thermal stability by assessing the apparent thermal melting temperature (T_m_) of the WT and stabilized derivatives through nano-DSF. We detected a notable increase (∼7°C) in the T_m_ of the stabilized derivatives relative to the wild type RBDs (Figure2C, S2A). To verify proper folding of these RBD derivatives, we examined the binding with a selected panel of broadly neutralizing antibodies (bNAbs) that bind to different epitopes on RBD by using SPR (Figure2D). Due to the lack of conservation in class 1 and class 2 RBD epitopes, antibodies within these categories typically lack significant cross-reactivity towards sarbecovirus RBDs. However, class 3 and class 4 antibodies bind to relatively conserved epitopes, thereby offering potential protection against emerging sarbecoviruses. The epitope of bnAb 10-40 is similar to the previously defined ‘class 4’ antibody epitope, it makes polar contacts and hydrophobic interactions with RBD residues (377-385). ADG-20 binds to a class ¼ epitope that overlaps with the ACE2 binding site. This bnAb binds to RBDs from clade 1a, 1b and 3; however there was no binding observed for the clade 2 RmYN02 RBD which also does not bind ACE2. S2X259 targets the conserved antigenic site II within the RBD. It interacts with amino acid residues 369-386, 404-411 and 499-508(36). All our RBD derivatives except the stabilized RmYN02 trimer bound to this bnAb, this could be due to loss of conservation of residue D405 and G504 in RmYN02 RBD which are crucial for interaction with S2X259. All the stabilized RBD derivatives also bound to class 4 bnAb CR3022 (Figure 2D, S2B-E).

**Figure 2:**
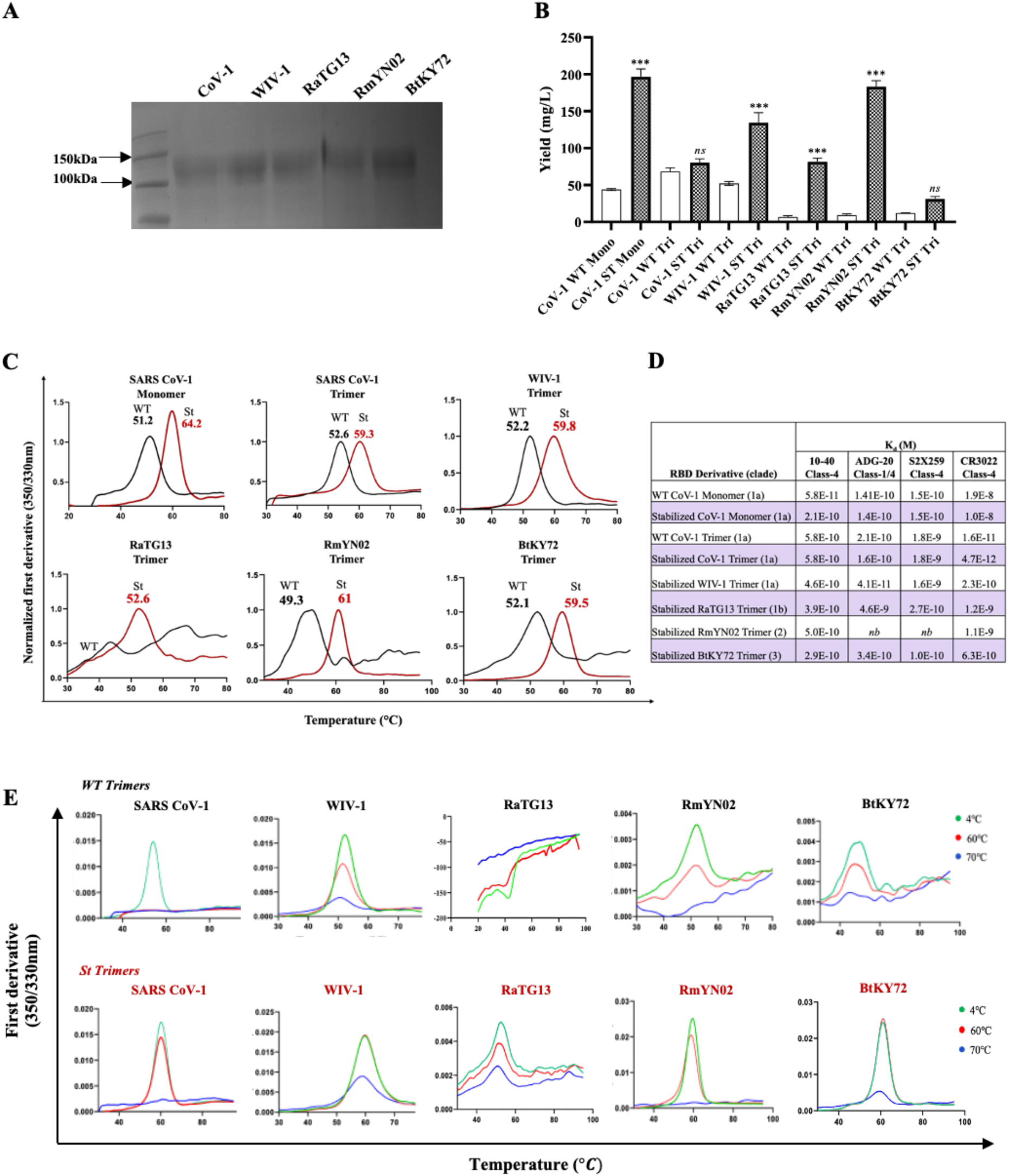
Biophysical characterization of sarbecovirus RBDs. (A) SDS-PAGE profile of stabilized trimeric sarbecovirus RBDs expressed in *Expi293F* cells (B) Yield of purified WT and stabilized sarbecovirus RBDs (C) Thermal denaturation profile of WT and stabilized RBDs (D) Binding affinity of sarbecovirus RBD derivatives with broadly neutralizing antibodies determined through SPR, *nb indicates no binding was observed* (E) Nano-DSF curves showing short-term thermal tolerance of RBD derivatives after incubation at the indicated temperatures for two hours. *p values were calculated by two-tailed Mann–Whitney test. *, **, and *** indicate p < 0.05, <0.01, and < 0.001, respectively*.

To probe the thermal tolerance conferred by these mutations, we incubated the WT and St RBD derivatives at different temperatures ranging from 4°C to 70°C for 2 hours. The proteins were then cooled to room temperature and subjected to DSF. The thermal melt curves clearly demonstrated the extended thermal tolerance of the stabilized derivatives relative to the corresponding WT RBDs (Figure 2E). Collectively, these results illustrate that indeed the identified mutations in SARS-CoV-2 RBD background are stabilizing in all the sarbecoviruses included in this study; they not only result in enhanced thermal stability and thermal tolerance but also increase the yield of the purified proteins significantly. Binding experiments confirm that the stabilized RBD derivatives are properly folded.

### Stabilized RBD derivatives and a corresponding cocktail formulation elicit neutralizing antibody responses in mice against diverse sarbecoviruses

To assess the immunogenicity of the stabilized sarbecovirus RBD derivatives, we immunized female C57BL/6 mice (n=5/group) intramuscularly with RBD derivatives (5μg each) adjuvanted with SWE in a prime-boost regimen. We evaluated the immunogenicity of a sarbecovirus RBD cocktail consisting of 5μg of each of the stabilized RBDs from SARS-CoV-1 and WIV-1 (clade 1a); SARS-CoV-2 and RaTG13 (clade 1b); RmYN02 (clade 2) and BtKY72 (clade 3) formulated with SWE as the adjuvant (Figure 3A). Post boost immunization, ELISA against respective RBDs was done to determine the serum IgG titres. High antibody titres were observed upon immunization with individual RBD derivatives, as well as with the RBD cocktail (Figure 3B). In the context of SARS-CoV-1 immunogens, there was considerable enhancement in the immunogenicity of the monomeric RBD derivative upon stabilization; however, in trimeric format this difference was much smaller (Figure 3C). The stabilized monomeric SARS-CoV-1 RBD derivative elicited higher neutralizing antibody (NAb) titres against homologous (SARS-CoV-1, WIV-1) and heterologous (LYRa3, SHC014) clade1a pseudoviruses related to WT RBD. In contrast, the WT and mutant trimeric derivatives elicited comparable neutralizing titres in most cases but the stabilized trimer elicited higher titres against SARS-CoV-1 (Figure 4A). Consistent with earlier data with WT SARS-CoV-2 RBD, WT SARS-CoV-1 RBD elicited considerably lower titers than stabilized RBDs (63). The RBD cocktail elicited good neutralization against homologous SARS-CoV-1, WIV-1 (clade 1a); SARS-CoV-2 B.1 (clade 1b) and BtKY72 (clade 3) as well as heterologous pseudoviruses (clade1a: LYRa3, SHC014; clade3: Khosta-2). Importantly with a single RBD immunization, clade matched pseudoviruses were effectively neutralized but no cross neutralization of mismatched pseudoviruses was observed (Figure 4 B-D, S3A-D). This emphasizes the need for incorporating RBDs from all the clades to achieve a broadly neutralizing pan-sarbecovirus response. Unfortunately, we could not assess neutralization titres against clade 2 viruses as they do not bind ACE2 and thus a suitable neutralization assay was not available. Overall, the RBD cocktail formulation elicited good neutralizing titres against all the RBDs incorporated in the formulation confirming the immunogenicity of our RBD derivatives in individual as well as cocktail format. It also elicited neutralization against all the heterologous pseudoviruses tested, indicating elicitation of a broad neutralization response in the immunized animals.

**Figure 3:**
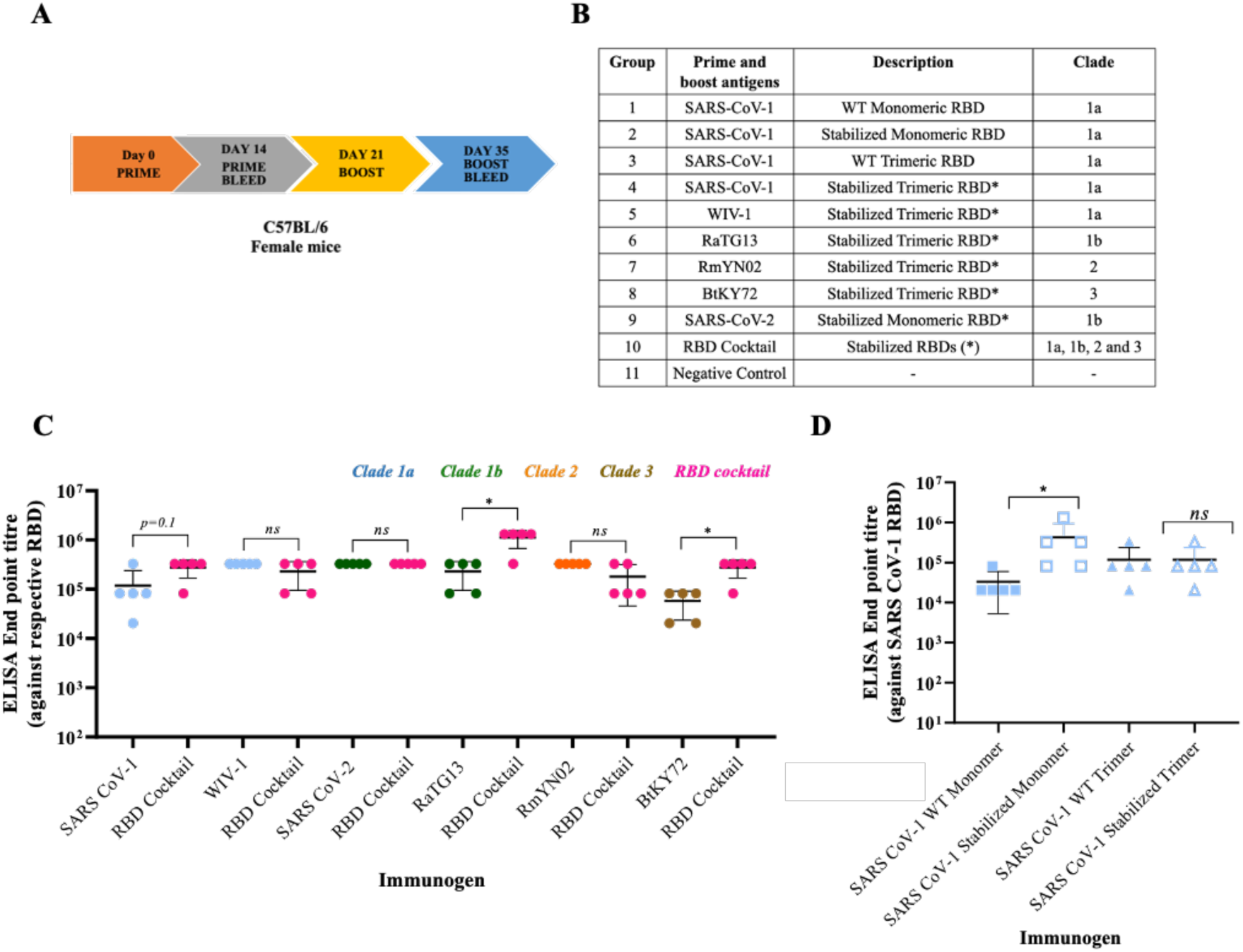
Immunogenicity of sarbecovirus RBDs in individual and cocktail formulations. (A) Schematic representation of the prime-boost immunization (B) Description of the sarbecovirus RBD derivatives used as individual immunogens and as components of the RBD cocktail (C) Comparison of ELISA end point titres of individual groups with the RBD cocktail against various RBDs (C) Comparison of IgG titres in groups immunized with SARS CoV-1 monomeric or trimeric RBDs with or without stabilizing mutations. *p values were calculated by two-tailed Mann–Whitney test. *, **, and *** indicate p < 0.05, <0.01, and < 0.001, respectively*.

**Figure 4:**
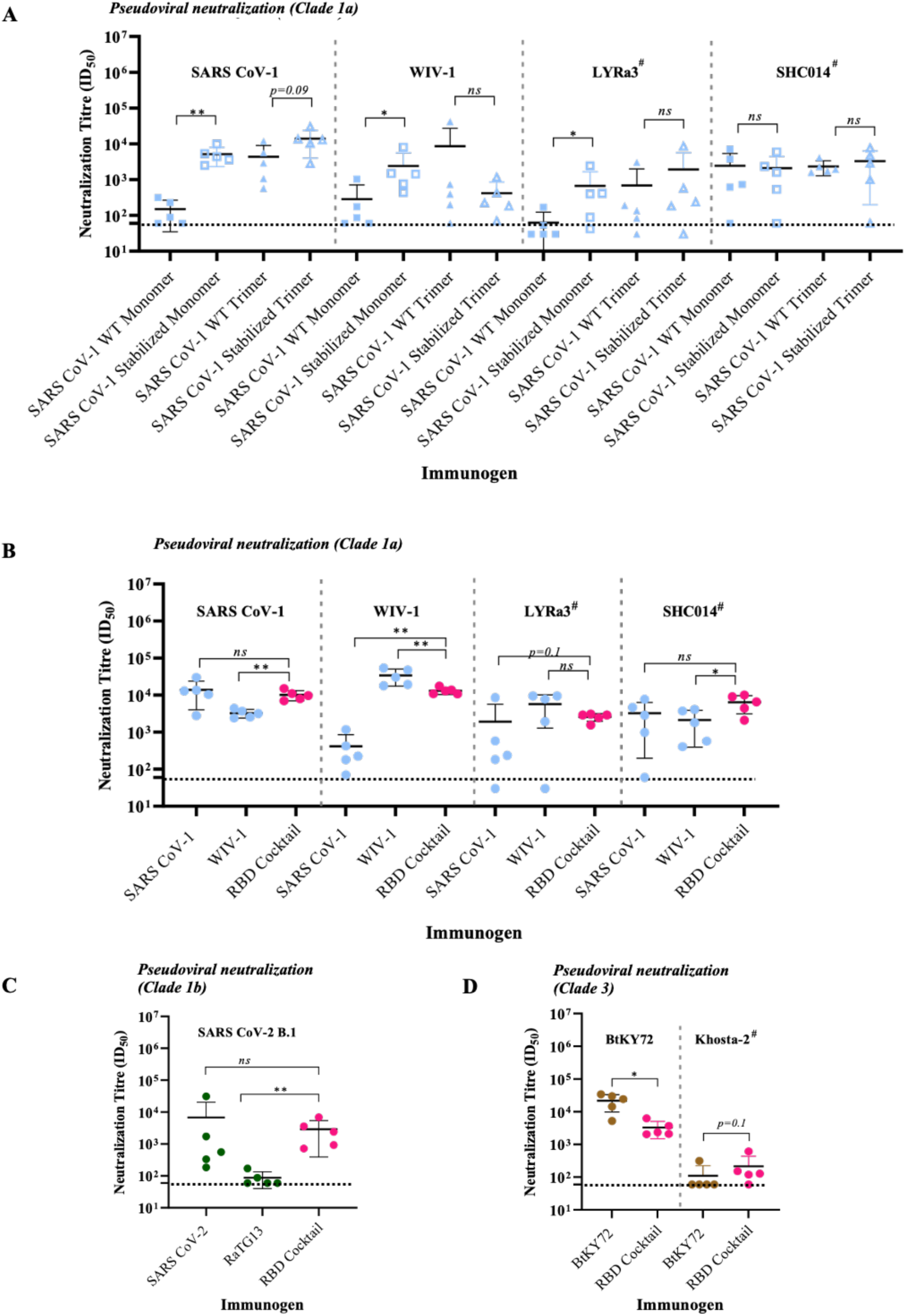
Homologous and heterologous pseudoviral neutralization titres elicited by stabilized RBD derivatives after two immunizations. (A) Neutralization of Clade 1a pseudo viruses by sera from groups immunized with SARS CoV-1 derived RBD immunogens. Pseudoviral neutralization titres against viruses from (B) Clade-1a (C) Clade-1b and (D) Clade-3 by clade specific single immunogen in comparison with the RBD cocktail group *[# represents heterologous pseudovirus]. p values were calculated by two-tailed Mann–Whitney test. *, **, and *** indicate p < 0.05, <0.01, and < 0.001, respectively*.

### Stabilized RBD cocktail induces an immunogenic response in pre-immunized mice

Currently, practically the entire human population has been exposed to SARS-CoV-2 through vaccination, infection or both. We therefore aimed to deduce whether the RBD cocktail formulation could induce comparable responses in pre-immunized animals and naïve mice. We therefore immunized female BALB/c mice (n=5/group) with SARS-CoV-2 B.1 RBD (5μg) followed by two boost immunizations with the stabilized RBD cocktail formulation (5μg of each antigen, total 30μg) (Figure 5 A). Serum ELISA demonstrated the presence of IgG titres against sarbecovirus RBDs after boost 1, boost 2 did not result in further increase in antibody titres (Figure 5B, S4A). However, with sera from the first boost, only weak and sporadic neutralization of homologous as well as heterologous pseudovirus was observed. Consistent, broad neutralization was observed only after the second boost of the RBD cocktail (Figure S4B-D). Consistent with the previous results in naïve mice; the pan-sarbecovirus formulation was able to elicit a broad neutralizing responses against sarbecoviruses after two immunizations in pre-immunized mice, confirming its relevance in the current scenario.

**Figure 5:**
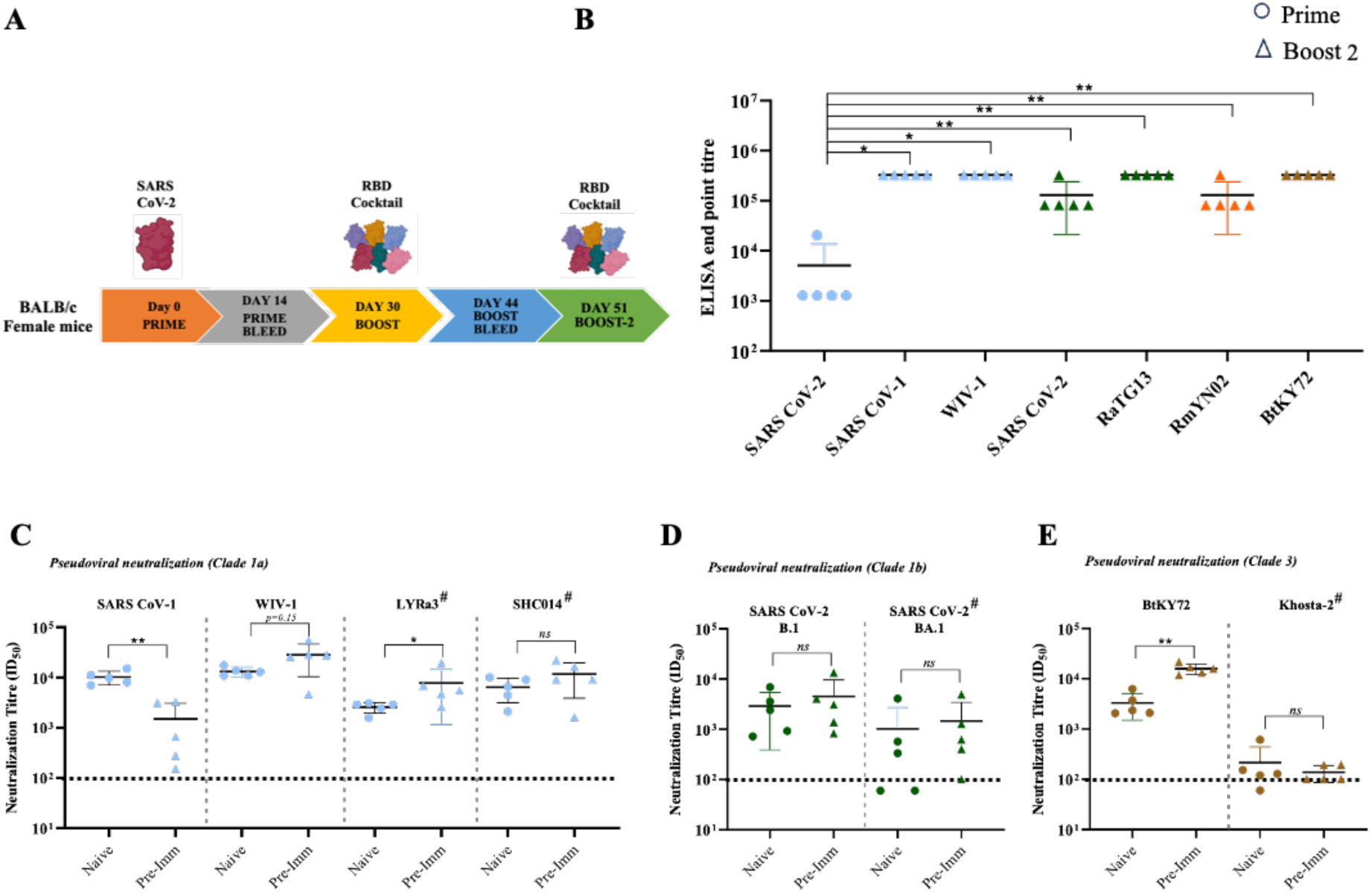
Immunogenicity of sarbecovirus RBD cocktail formulation in pre-immunized mice. (A) Immunization scheme in mice, consisting of prime with stabilized SARS CoV-2 RBD and two boost immunizations with RBD cocktail formulation (B) ELISA end point titres against SARS CoV-2 RBD after prime immunization and sarbecovirus RBDs after boost immunization. Neutralization of (C) clade-1a, (D) clade-1b and (E) clade-3 pseudoviruses after a second boost immunization of pre-immunized mice relative to naïve immunized group *[data for pseudoviral neutralization after boost in naïve mice is taken from Figure4; # represents heterologous pseudovirus]*. *p values were calculated by two-tailed Mann–Whitney test. *, **, and *** indicate p < 0.05, <0.01, and < 0.001, respectively*.

### A lyophilized RBD cocktail retains antigenicity for upto a month at 37°C

To assess the long-term stability of the RBD cocktail formulation, we subjected it to lyophilization and incubated the lyophilized material at either 4 or 37°C for a duration of one month. The stability of our formulation was confirmed through DSF and BLI experiments. As depicted in Figure 6A, the lyophilized formulation stored at both 4 and 37°C exhibited similar melting profiles until day 15. However, a slight reduction in stability was observed on day 30. Additionally, we assessed the conformational integrity of the RBD cocktail by examining binding with previously described bNAbs (CR3022, 10-40, ADG-20, S2X259). Consistent with the DSF analysis, the lyophilized formulation demonstrated appropriate binding curves with all the antibodies until day 15, with a slight decrease in binding signal observed on day 30 (Figure 6B). These findings demonstrate the long-term thermal stability of the cocktail formulation when in a lyophilized state.

**Figure 6:**
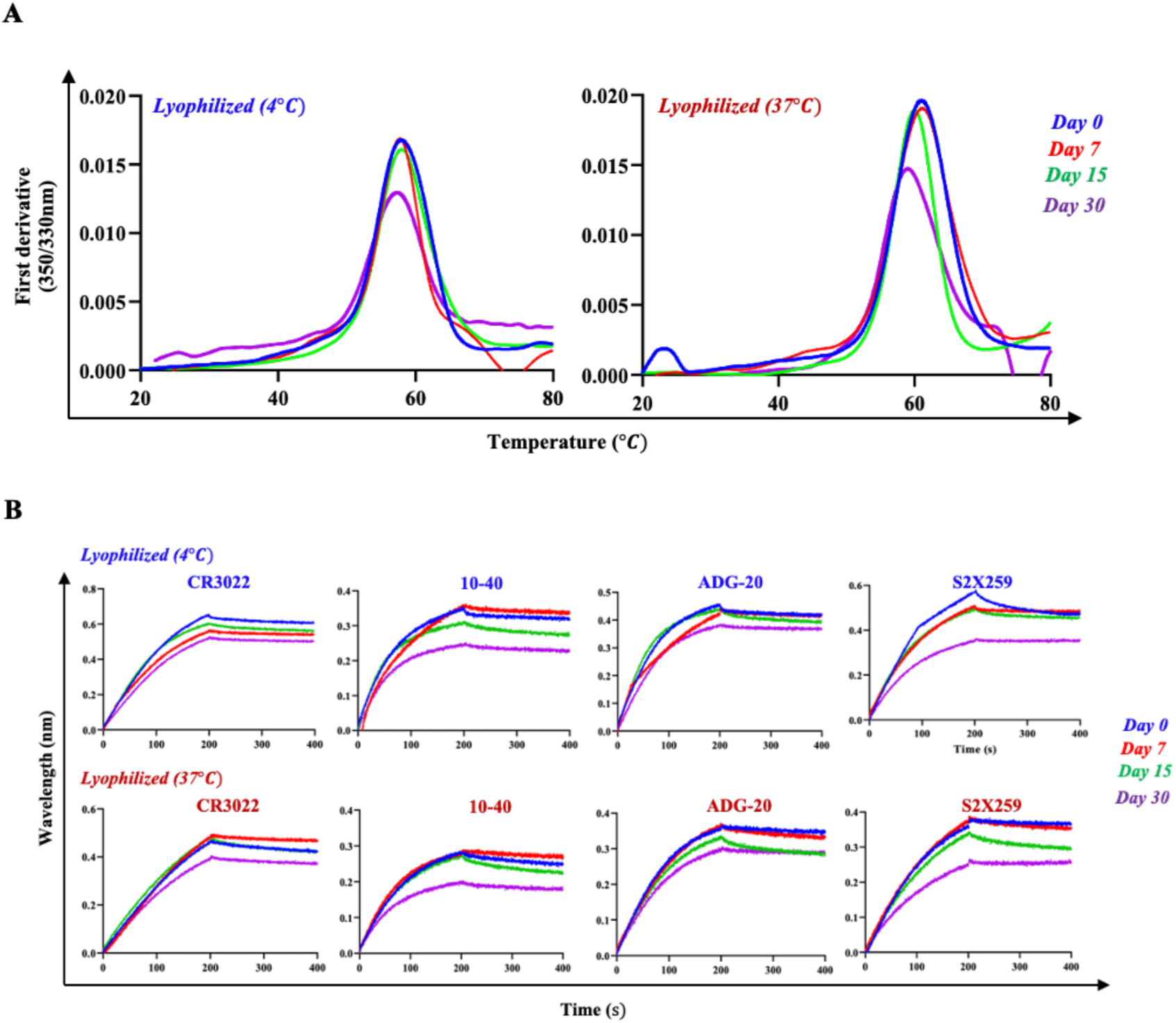
Long-term thermal stability of the RBD cocktail immunogens. (A) Thermal denaturation profile and (B) binding affinity of the RBD cocktail immunogens from day0-day 30 for proteins incubated at 4°C and 37°C determined by Nano-DSF and BLI experiments respectively. Lyophilized proteins were incubated at the indicated temperatures and reconstituted in PBS prior to measurements.

### Structure of stabilized SARS-CoV-2 RBD

To analyse whether the three mutations-A348P, Y365W, and P527L in RBD have any effect on the overall architecture of the RBD, we performed a single particle cryo-EM study of a SARS-CoV-2 spike containing the mutations. Our final cryo-EM map at 4.6 Å resolution demonstrated a compact architecture of the three RBDs along with NTD and S2 domains, similar to our previous study with the S2P spike ectodomain(57) (Figure 7A). We saw densities corresponding to all three RBDs in our cryo-EM map (Figure 7A). Furthermore, we observed an overall good fitting of the real space refined model in the cryo-EM density, suggesting that the resultant model was suitable for further analysis (Figure 7B). To ascertain if the mutations induced any local rearrangements in the RBD, we aligned the wildtype S protein atomic model (PDBID: 6VXX) with our real-space refined mutant S protein model from the current study (Figure 7C). There were no differences in the overall architecture between the wildtype and stabilized S ectodomains. Interestingly, the three mutations did not induce any significant local structural rearrangements in the RBD, and the overall RBD fold was maintained (Figure 7C). This confirmed that the mutations-A348P, Y365W and P527L do not disrupt the overall fold of RBD. Although we did not observe significant densities for P348 and L527, we obtained strong density corresponding to W365 (Figure 7D). W365 engages with neighbouring residues, L387 and F515 with distances of 3.6 and 3.7Å respectively (Figure 7E), thus contributing to the local stability of the RBD fold. We also assessed the existing RBD structures for the sarbecoviruses mentioned in this study and found that the conserved proline residue (corresponding to the A348P mutation identified in SARS-CoV-2) interacts with either Gln or Glu six residues downstream. Proline is known to form C-H-O weak non bonded interactions(58). In addition, proline stabilizes proteins through reduction of the conformational entropy of the denatured state(59,60). In both RaTG13 and SARS-CoV2 (clade 1b sarbecoviruses), this proline is replaced by alanine, abrogating this interaction. With A348P mutation, the proline is now positioned to interact with the Asn354 residue (six residues downstream), providing additional stability to the local architecture similar to other sarbecoviruses (Figure 7F). Overall, our cryo-EM study reports that these mutations do not induce any major structural rearrangements, maintains the overall structure of RBD as compared to wildtype construct and provides structural insights into the origins of the enhanced stability arising from the mutations.

**Figure 7:**
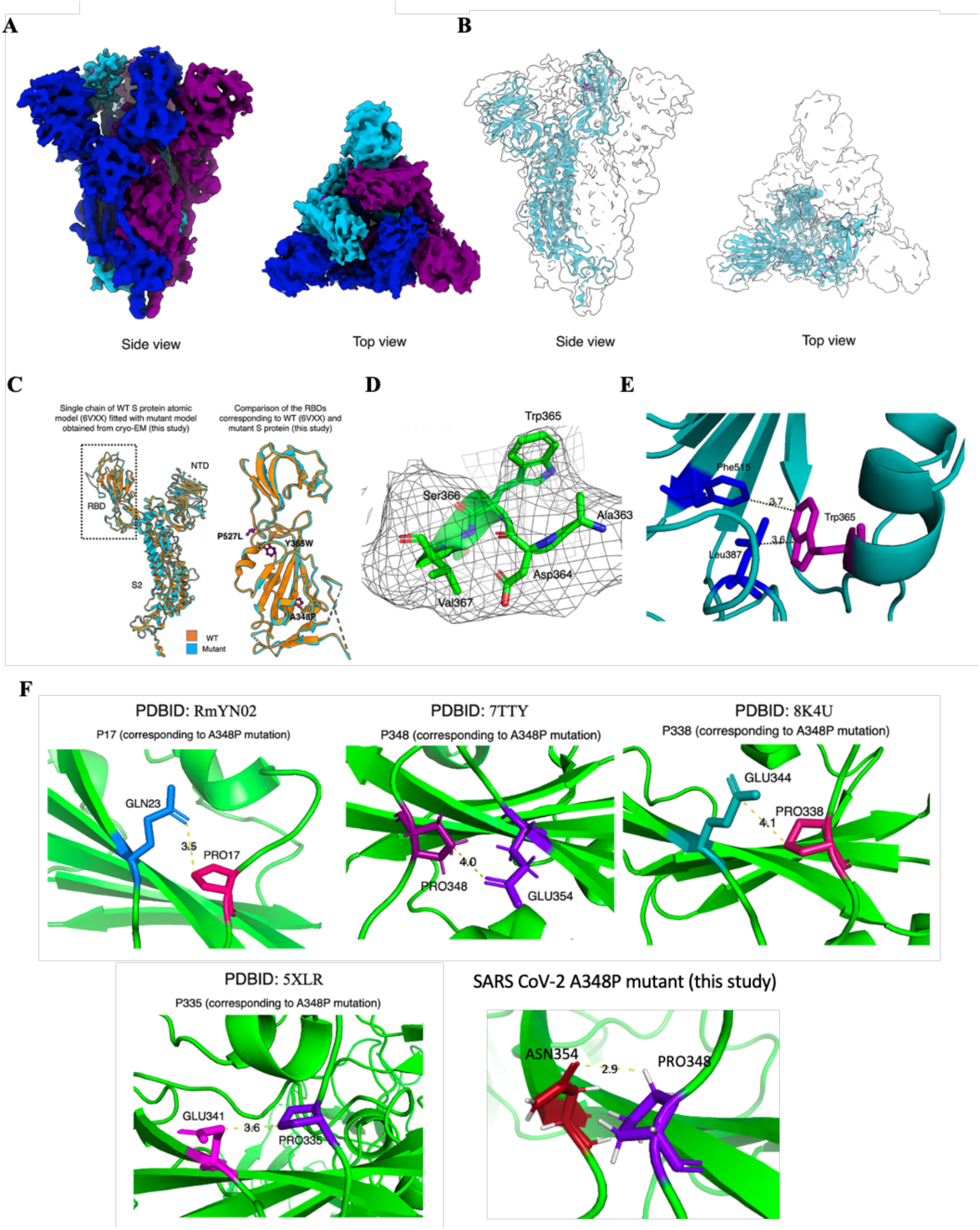
Effect of mutations on RBD conformation. (A) Side view (left) and top view (right) of the cryo-EM map of mutant SARS CoV-2 spike protein. The three monomers are marked in different colors (dark blue, dodger blue, and purple). (B) Fitting of the atomic model (dodger blue) is shown for one monomer in the translucent cryo-EM map for clarity. (C) Alignment of real space refined model from this study (dodger blue) with wildtype S protein (PDBID: 6VXX, orange) is shown. Zoomed in view of the RBD region is provided on the right. (D) Cryo-EM density corresponding to the mutation Y365W in mesh view at sigma level 5. Atomic model is colored in green. (E) Neighboring residues are shown for W365, and their distances are shown with dotted lines. Numeric values of distance are in Å. (F) Neighboring interacting residues shown for P348 for RmYN02 (modelled structure), WIV-1 (PDBID: 7TTY), BtKY72 (PDBID: 8K4U), SARS CoV-1 (PDBID: 5XLR), and SARS CoV-2 (this study).

## Discussion

Sarbecoviruses are a subgenus of betacoronaviruses that includes the severe acute respiratory syndrome coronavirus (SARS-CoV), Middle East respiratory syndrome coronavirus (MERS-CoV), and the more recently identified SARS-CoV-2, which causes COVID-19(5,9,11). The potential for future pandemics is associated with the capacity of a virus to infect humans, cause sustained human-to-human transmission, and evade pre-existing immunity. Several sarbecoviruses closely related to SARS-CoV-2 have been identified in bats, the presumed natural reservoir. These include WIV-1, RaTG13, RmYN02 and BtKY72 among many others(52–56). While they may not currently pose an immediate threat, their potential for spillover into humans raises concerns, especially given the experience with SARS-CoV-1, MERS-CoV and SARS-CoV-2. Recent studies have employed varied approaches to develop broadly neutralizing vaccine candidates ranging from multi-antigen DNA(61), mRNA to mosaic nanoparticle platforms(43–45,47,48). Immunization studies with mosaic nanoparticles composed of heterotypic RBDs from different sarbecoviruses have shown promising results in mice and NHPs(43). Another study which utilized a trivalent nanoparticle approach based on sarbecovirus as well as merbecovirus RBD demonstrated protection against SARS-CoV and MERS-CoV in mice(46). In most prior studies, however protein yields and thermal tolerance of the antigens was not reported. The stabilized RBD cocktail formulation reported here elicits high antibody binding response and pseudovirus neutralization titres in mice, however we were unable to assess protective efficacy due to unavailability of authentic sarbecoviruses.

The receptor binding domain of the SARS-CoV-2 spike protein has emerged as a potent and versatile vaccine candidate with distinct advantages. Its ability to induce a robust immune response, specificity, potential for cross-protection, and practical considerations for large-scale production collectively position the RBD as an important molecule in the global effort to combat future coronavirus pandemics(28–31). Previous reports from our group demonstrated the stabilization of SARS-CoV-2 RBD which resulted in enhanced thermostability, lower dynamic flexibility as well as higher immunogenicity(28–30,62) including against SARS-CoV-2 Omicron lineage variants (63). Interestingly, A348P is a naturally occurring substitution on Clade 1a, 2 and 3 sarbecoviruses, it is probable that this substitution might play a role in enhancing protein stability by reducing the conformational entropy of the unfolded state, consequently increasing the free energy of the unfolded state. Y365W is a cavity filling mutation while the origin of the stabilizing impact of the P527L substitution is currently unclear, due to the disordered nature of this residue in the crystal structure of RBD when bound to ACE2.

In this study, we demonstrate the transferability of these stabilizing mutations identified in SARS-CoV-2 RBD to diverse sarbecovirus RBDs having varying similarity with SARS-CoV-2 (Figure 1B-D); suggesting these mutations could potentially be effective for other clade members as well. We have expressed the stabilized trimeric RBD derivatives in mammalian cells and reported high yields of purified proteins (Figure 2A, B). The stabilized RBD derivatives showed significant enhancement in the apparent melting temperature and short-term thermal tolerance relative to corresponding WT RBDs (Figure 2C-E). All the stabilized RBDs elicited high IgG titres in mice in individual and cocktail format (Figure 3) and thus proved to be highly immunogenic. The RBD cocktail formulation performed comparable or better than the individual clade-matched ability to elicit homologous and heterologous pseudoviral neutralization respectively (Figure 4). Overall, the absence of cross-reactivity within clades appears to be evident based on the findings from pseudoviral neutralization assays following vaccination with individual RBD derivatives (Figure S3). This underscores the importance of exposure to a variety of antigens in order to develop a broad response, such as the cocktail formulation that we employed. We observed that in addition to naive mice, our RBD cocktail formulation is immunogenic in SARS-CoV-2 RBD pre-immunized mice as well (Figure 5), mirroring the present situation after COVID-19 vaccination and infection. While in the present study we have focused on antibody responses, we have previously shown (63) that a similar monovalent formulation of the stabilized SARS-CoV-2 RBD elicits a balanced T-cell response in mice. Further, the RBD cocktail immunogens exhibited good thermal stability following lyophilization for a period of up to two weeks when stored at 37°C (Figure 6) and we have previously shown for the monovalent SARS-CoV-2 formulation that immunogenicity is also retained after lyophilization, storage for one month at 37°C, and subsequent resolubilization and immunization (64). During the COVID-19 pandemic, adenoviral vector and mRNA-based vaccine formulations were the most rapidly developed and widely used in LMICs and developed countries respectively. The former were less immunogenic but were cheaper and could be stored under refrigerated conditions. However, there were concerns with associated side effects (65,66), and anti-vector immunity upon repeated immunization may also be an issue(67–70). mRNA vaccines were more expensive and had stringent low temperature requirements. The present protein subunit formulation overcomes some of these issues. In summary, we show that stabilization and production of diverse sarbecovirus RBD derivatives to be used as vaccine antigens is technically feasible. The significant improvement in yield and thermostability of immunogens by transfer of identified stabilizing mutations as described, would enhance the manufacturing efficiency and distribution of RBD-based vaccines for sarbecovirus on a global scale, addressing the pressing need for quick, widespread and equitable vaccination in case of future sarbecovirus epidemics and pandemics.

## Methods

### Protein Expression and Purification

The genes encoding sarbecovirus RBD proteins and monoclonal antibodies (mAbs) were synthesized at GenScript (USA) and TWIST Biosciences (USA). All the RBD derivatives were purified from transiently transfected *Expi293F* cells following the manufacturer’s guidelines (Gibco, Thermo Fisher, Waltham, MA, USA). Briefly, *Expi293F* cells were diluted to a density of 3 × 10^6^ cells/ml. For transfection, the desired plasmid was complexed with ExpiFectamine293 according to the manufacturer’s protocol and transiently transfected into Expi293F cells. Enhancer 1 and Enhancer 2 were added 16h post transfection. The culture supernatant was collected after 5 days, and protein was purified through Ni-NTA affinity chromatography using Ni Sepharose 6 fast-flow resin (GE Healthcare, Chicago, IL, USA) for RBD derivatives. The supernatant was added to a pre-equilibrated Ni-NTA column. Following a 2 column wash with 1× PBS (pH 7.4) supplemented with 20 mM imidazole, the protein was eluted in 1X PBS with 300mM imidazole (pH 7.4). Pooled eluted fractions were dialyzed thrice against 1× PBS (pH 7.4). For expression and purification of mAbs, Expi293F cells were co-transfected by Heavy and Light chain plasmids in 1:1 ratio by using polyethylenimine, and supernatants were harvested after 5 days post-transfection. The antibodies were purified from supernatants by using Protein A/G beads, dialyzed and stored in phosphate buffered saline (PBS) for further use. Purified protein samples were analyzed on 12% SDS-PAGE gel and quantified using NanoDrop spectrophotometer.

### Thermal Unfolding experiments

Thermal melting studies of the RBD derivatives were performed using nanoDSF (Prometheus NT.48), as described(71) at a temperature range of 20 °C to 95 °C. Experimental measurements were conducted at 100% LED intensity with an initial discovery scan, and the scan counts (350 nm) ranged between 2000 and 3000. For lyophilized protein reconstitution was done using 1X PBS buffer before the DSF experiment.

### SPR binding studies

Kinetics titrations were performed using a CM5 sensor chip (Cytiva) at 25°C. The activation of the carboxymethylated-dextran gold surface was achieved by injecting EDC/s-NHS (2/4 mM) solution in autoclaved water pH 7.0 injected at 5 µL/min for 400 seconds. Following the activation step, a 10 µg/mL solution of Protein-G in sodium acetate (NaOAc) pH 5.0 was injected over the activated surface at 5 µL/min. After covalent modification of the sensor surface, a quenching solution of ethanolamine pH 8.5 (Cytiva) was injected over the surface for 600 seconds to cap any residual active NHS esters. PBS (1X) pH 7.4 was used for the running buffer during titration. During the kinetics assay, one flow cell channel with only Protein G served as a reference channel to monitor and subtract binding responses due to non-specific interactions. 200-300 RU of monoclonal antibodies at 1 µg/mL were captured onto the chip surface for each cycle at 5 µL/min for 60 seconds, followed by injection of RBD derivatives for 200 seconds. Then a dissociation step was performed using an injection of running buffer for 200 seconds. Following the dissociation step, regeneration of the Protein G surface was performed using 1 injection of 0.1M glycine-HCl, pH 2.0 at 30 µL/min for 40 seconds. The flow rate for association and dissociation was 30 µL/min. The kinetics traces were reference subtracted using the responses of the reference channel in each cycle and blank subtracted using a zero-concentration cycle. Then the kinetics constants k_a_, k_d_ and K_D_ values were determined using Biacore T200 evaluation software.

### Mice Immunizations

Female C57BL/6 mice (6–8 weeks old, n = 5/group) were immunized intramuscularly with RBD derivatives (5µg/animal in 100 µL of 1× PBS, pH 7.4) adjuvanted with SWE (1:1 v/v antigen: adjuvant ratio) (Sepivac SWE Batch No. 200915012131, Cat. No. 80748J, SEPPIC SA) on days 0 (prime), and 21 (boost). RBD cocktail group was immunized with 5µg/antigen having RBD derivatives from different clades as mentioned. Sera were isolated from blood drawn on days prior to prime (day −1), post-prime (day 14), and post-boost (day 35). After completion of the experiment, the animals were euthanized with overdose of isoflurane (5%) followed by cervical dislocation.

For the assessment of RBD cocktail efficacy in pre-immunized mice, female BALB/c mice (6–8 weeks old, n = 5/group) were first immunized intramuscularly with stabilized monomeric SARS-CoV-2 RBD (1µg/animal in 100 µL of 1× PBS, pH 7.4) adjuvanted with SWE (1:1 v/v antigen: adjuvant ratio) (Sepivac SWE Batch No. 200915012131, Cat. No. 80748J, SEPPIC SA) on days 0 (prime), and then boosted with RBD cocktail formulation (5µg/antigen, total 30µg) on day 30 and day 51. Previous COVID-19 vaccine studies have used both BALB/c and C57BL/6 mice with similar results(72,73). Sera obtained before prime and post boost immunizations was used to conduct ELISA and pseudoviral neutralization assays. Blood was collected through retro-orbital puncture by anaesthetizing the animals with 1-2% of isoflurane briefly for ∼10 seconds. After completion of the experiment, the animals were euthanized with overdose of isoflurane (5%) followed by cervical dislocation. These studies were performed at Central Animal Facility, Indian Institute of Science. The Institutional Animal Ethics committee approved all animal studies (IAEC no. CAF/ETHICS/002/2023).

### Enzyme-linked immunosorbent assay (ELISA)

ELISA was used to quantify the sarbecovirus RBD-specific IgG titers in serum as described previously(28). Briefly, ELISA plates were coated with 4 µg/mL His tagged RBD derivatives at 25 °C, 1h. The coated plates were blocked using blocking buffer (PBS containing 3% skimmed milk) for 1 h at 25 °C. The sera (maximum concentration, 1:80-1:1280) from mice, were serially diluted 1:4 in PBST (PBS containing 3% skimmed milk and 0.05% Tween-20) for 8 dilutions, followed by incubation with the coated ELISA plates for 60 min at 25 °C. After washing with PBST three times, HRP-conjugated secondary antibody was added to the plates and incubated for 60 min at 25 °C. After incubation with the secondary antibody, the plates were washed with PBST three times, followed by adding 3,3’,5,5’-tetramethylbenzidine (TMB) to visualize the reaction. Finally, 6N HCl was used to stop the reaction. The chromogenic signal was measured at 405 nm using an ELISA plate reader (Maxome Labsciences Cat # P3-5x10NO). The serum dilution with a signal observed two-fold above the negative control (empty blocked wells) was considered the endpoint titer for ELISA.

### Sarbecovirus Pseudovirus Preparation and Neutralization Assay

HIV-1-based pseudotyped viruses were employed in pseudoviral neutralization assays, following a method previously described(29). Briefly, adherent HEK293T cells were transiently transfected with plasmid DNA pHIV-1 NL4-3Δenv-Luc and sarbecovirus spike plasmids, using the ProFection mammalian transfection kit (Cat# E1200, Promega Inc., Singapore) for pseudovirus production. The genes encoding Spike proteins from SARS-CoV-2 VOCs were synthesized at GenScript (USA) while spike plasmids encoding sarbecovirus spikes were kindly gifted by Dr. Pamela J. Bjorkman (Caltech, USA). The culture supernatant was harvested 48 h post-transfection, filtered through a 0.22 μm filter, and stored at −80 °C. Adherent HEK293 cells expressing hACE-2 and TMPRSS2 receptors (BEI resources, NIH, Catalog No. NR-55293) were cultured in a growth medium consisting of DMEM with 5% Fetal Bovine Serum (Thermo Fisher) and penicillin–streptomycin (100 U/mL). Mice serum samples were heat-inactivated and then serially diluted in the growth medium, starting from 1:20 dilutions. In the next step, the pseudotyped virus was incubated with the serially diluted sera in a total volume of 100 µL for 1 h at 37 °C. The adherent cells were then trypsinized, and 1 × 104 cells/well were added to achieve a final volume of 200 µL/well. The plates were further incubated for 48 h in a humidified CO_2_ incubator at 37 °C. After incubation, neutralization was measured as an indicator of luciferase activity in the cells (relative luminescence units) using Nano-Glo luciferase substrate (Cat # N1110, Promega). Luminescence was measured using a Cytation-5 multimode reader (Bio-Tech Inc., Oklahoma City, OK, USA). The luciferase activity, measured as relative luminescence units (RLU), in the absence of sera was considered as 100% infection. The serum dilution resulting in half-maximal neutralization of the pseudovirus (ID50) relative to the no-serum control was determined from neutralization curves.

### Biolayer Interferometry experiment

The long term stability of lyophilized RBD cocktail was assessed through DSF and BLI binding studies. BLI measurements were made using ForteBio biosensors (Fortebio - Sartorius). All data collection were performed at 25°C using settings of Standard Kinetics Acquisition rate at a sample plate shake speed of 1000 rpm. mAbs were loaded onto Protein G sensors, subsequently they were dipped into 1x PBS buffer for 60 seconds to obtain baseline and then dipped into wells containing RBD derivatives at different concentrations in 1X PBS to monitor antibody association. The dissociation step was monitored for 200 seconds by dipping Ab-bound sensors into buffer. Antigen specific binding responses were obtained by subtracting responses of blank sensors tested in parallel with 1X kinetics buffer. The specific binding responses were fitted using ForteBio Data Analysis 12.0 software tool.

### Cryo-EM sample preparation and data acquisition

Prior to vitrification, R1.2/1.3 300 mesh gold grids (Quantifoil, Electron Microscopy Sciences) were glow-discharged at 20 mA for 130 sec. 3 μl freshly prepared sample was added to the grid and incubated for 10 sec before blotting for 7 sec at 100 percent humidity, followed by rapid plunge freezing in liquid ethane using FEI Vitrobot Mark IV plunger. Data acquisition was performed with Talos Arctica (Thermo Scientific) at 200 kV, equipped with K2 Summit Direct Electron Detector. Latitude S (Gatan) was used for automatic data collection at nominal magnification of 54000x with pixel size 0.92 Å at specimen level, with defocus range of -0.75 and -2.25 μm and calibrated dose of 2-4 e-/ Å^2^ per frame for a total of 20 frames recorded over 8 sec. A total of 3239 movies were acquired for mutant S protein for image processing.

### Cryo-EM image processing pipeline

Raw movies were motion-corrected in RELION using MotionCorr2 and CTF was estimated on the resultant integrated image files. About 2000 particles were manually picked and classified, good classes corresponding to different orientations were further used for template-based automatic picking. With a box size of 336 pixels, particles were obtained and subjected to rigorous 2D classification. Finally, 533492 particles were further used for two rounds of 3D classification. All resultant classes from the second 3D classification were refined individually and in different combinations, and the final structure was refined with C3 symmetry using a total of 388200 particles, which yielded the map with the highest resolution and sharpest features. Resolution was estimated based on GS-FSC. PHENIX was used for rigid body docking followed by real space refinement. Visualization was performed using ChimeraX and PyMOL.

### Statistical Analysis

The p values for ELISA binding and neutralization titres were analysed with a two-tailed Mann–Whitney test using the GraphPad Prism software 9.0.0 (* indicates p < 0.05, ** indicates p < 0.01, **** indicates p < 0.0001). VOC pseudoviral neutralization titre data were analysed with non-parametric Kruskal–Wallis with Dunn’s multiple-comparison tests using the GraphPad Prism software 9.0.0 (* indicates p < 0.05, ** indicates p < 0.01, **** indicates p < 0.0001).

## Data availability

All data are available in the main text or the supplementary materials.

## Acknowledgements

This work was supported by the Bill and Melinda Gates Foundation grant (INV-042471). SS acknowledges the Prime Minister Research Fellowship for her fellowship (PM/MHRD-20-16691.03). R.V. is a JC Bose Fellow of DST. S. K. thanks CSIR for senior research fellowship. We acknowledge DST FIST, MHRD, and the DBT IISc Partnership Program for infrastructural support. RPR acknowledges funding support from SERB (IPA/2020/000168) and fellowship from DBT. We acknowledge Prof. P. Bjorkman for providing the sarbecovirus spike constructs. The funders had no role in study design, data collection and interpretation, or the decision to submit the work for publication.

## Contributions

Conceptualization: R.V., S.S. Methodology: R.V., S.S., S.K., S.M., R.S.R., R.S., S.D., R.P.R. Investigation S.S., S.K., S.M., R.S.R., R.S. Visualization: R.V., S.S., S.K., S.M., R.S.R., R.S., S.D., R.P.R. Funding acquisition: R.V. Project administration: R.V. Supervision: R.V. Writing – original draft: S.S. Writing – review & editing: All authors.

## References

1. Ar Gouilh M, Puechmaille SJ, Diancourt L, Vandenbogaert M, Serra-Cobo J, Lopez Roïg M, et al. SARS-CoV related Betacoronavirus and diverse Alphacoronavirus members found in western old-world. Virology. 2018 Apr;517:88–97.

2. Temmam S, Vongphayloth K, Baquero E, Munier S, Bonomi M, Regnault B, et al. Bat coronaviruses related to SARS-CoV-2 and infectious for human cells. Nature. 2022 Apr;604(7905):330–6.

3. Evans TS, Tan CW, Aung O, Phyu S, Lin H, Coffey LL, et al. Exposure to diverse sarbecoviruses indicates frequent zoonotic spillover in human communities interacting with wildlife. Int J Infect Dis. 2023 Jun;131:57–64.

4. Gilbert M, Mohamed M, Choong SS, Baqi A, Kumaran J V, Sani I, et al. Presence of SARS-CoV-2-like coronaviruses in bats from east coast Malaysia. Trop Biomed. 2023 Sep 1;40(3):273–80.

5. Li Q, Guan X, Wu P, Wang X, Zhou L, Tong Y, et al. Early Transmission Dynamics in Wuhan, China, of Novel Coronavirus-Infected Pneumonia. N Engl J Med. 2020 Mar 26;382(13):1199–207.

6. Zhong NS, Zheng BJ, Li YM, Poon, Xie ZH, Chan KH, et al. Epidemiology and cause of severe acute respiratory syndrome (SARS) in Guangdong, People’s Republic of China, in February, 2003. Lancet. 2003 Oct 25;362(9393):1353–8.

7. Drosten C, Günther S, Preiser W, van der Werf S, Brodt HR, Becker S, et al. Identification of a novel coronavirus in patients with severe acute respiratory syndrome. N Engl J Med. 2003 May 15;348(20):1967–76.

8. Ksiazek TG, Erdman D, Goldsmith CS, Zaki SR, Peret T, Emery S, et al. A novel coronavirus associated with severe acute respiratory syndrome. N Engl J Med. 2003 May 15;348(20):1953–66.

9. Peiris JSM, Lai ST, Poon LLM, Guan Y, Yam LYC, Lim W, et al. Coronavirus as a possible cause of severe acute respiratory syndrome. Lancet. 2003 Apr 19;361(9366):1319–25.

10. Hijawi B, Abdallat M, Sayaydeh A, Alqasrawi S, Haddadin A, Jaarour N, et al. Novel coronavirus infections in Jordan, April 2012: epidemiological findings from a retrospective investigation. East Mediterr Health J. 2013;19 Suppl 1:S12–8.

11. Zaki AM, van Boheemen S, Bestebroer TM, Osterhaus ADME, Fouchier RAM. Isolation of a novel coronavirus from a man with pneumonia in Saudi Arabia. N Engl J Med. 2012 Nov 8;367(19):1814–20.

12. Zhou Z, Zhu Y, Chu M. Role of COVID-19 Vaccines in SARS-CoV-2 Variants. Front Immunol. 2022;13:898192.

13. Fiolet T, Kherabi Y, MacDonald CJ, Ghosn J, Peiffer-Smadja N. Comparing COVID-19 vaccines for their characteristics, efficacy and effectiveness against SARS-CoV-2 and variants of concern: a narrative review. Clin Microbiol Infect. 2022 Feb;28(2):202–21.

14. Walsh EE, Frenck RW, Falsey AR, Kitchin N, Absalon J, Gurtman A, et al. Safety and Immunogenicity of Two RNA-Based Covid-19 Vaccine Candidates. N Engl J Med. 2020 Dec 17;383(25):2439–50.

15. Jackson LA, Anderson EJ, Rouphael NG, Roberts PC, Makhene M, Coler RN, et al. An mRNA Vaccine against SARS-CoV-2 - Preliminary Report. N Engl J Med. 2020 Nov 12;383(20):1920–31.

16. Mathieu E, Ritchie H, Ortiz-Ospina E, Roser M, Hasell J, Appel C, et al. A global database of COVID-19 vaccinations. Nat Hum Behav. 2021 Jul;5(7):947–53.

17. Hare J, Hesselink R, Bongers A, Blakeley P, Riggall G. Improving vaccine equity by increasing vaccine thermostability. Sci Transl Med. 2024 Jul 24;16(757):eadm7471.

18. Lan J, Ge J, Yu J, Shan S, Zhou H, Fan S, et al. Structure of the SARS-CoV-2 spike receptor-binding domain bound to the ACE2 receptor. Nature. 2020 May;581(7807):215–20.

19. Tai W, He L, Zhang X, Pu J, Voronin D, Jiang S, et al. Characterization of the receptor-binding domain (RBD) of 2019 novel coronavirus: implication for development of RBD protein as a viral attachment inhibitor and vaccine. Cell Mol Immunol. 2020 Jun;17(6):613–20.

20. Hoffmann M, Kleine-Weber H, Schroeder S, Krüger N, Herrler T, Erichsen S, et al. SARS-CoV-2 Cell Entry Depends on ACE2 and TMPRSS2 and Is Blocked by a Clinically Proven Protease Inhibitor. Cell. 2020 Apr 16;181(2):271–280.e8.

21. Rogers TF, Zhao F, Huang D, Beutler N, Burns A, He WT, et al. Isolation of potent SARS-CoV-2 neutralizing antibodies and protection from disease in a small animal model. Science. 2020 Aug 21;369(6506):956–63.

22. Brouwer PJM, Caniels TG, van der Straten K, Snitselaar JL, Aldon Y, Bangaru S, et al. Potent neutralizing antibodies from COVID-19 patients define multiple targets of vulnerability. Science. 2020 Aug 7;369(6504):643–50.

23. Pinto D, Park YJ, Beltramello M, Walls AC, Tortorici MA, Bianchi S, et al. Cross-neutralization of SARS-CoV-2 by a human monoclonal SARS-CoV antibody. Nature. 2020 Jul;583(7815):290–5.

24. Huo J, Zhao Y, Ren J, Zhou D, Duyvesteyn HME, Ginn HM, et al. Neutralization of SARS-CoV-2 by Destruction of the Prefusion Spike. Cell Host Microbe. 2020 Sep 9;28(3):445–454.e6.

25. Wu Y, Wang F, Shen C, Peng W, Li D, Zhao C, et al. A noncompeting pair of human neutralizing antibodies block COVID-19 virus binding to its receptor ACE2. Science. 2020 Jun 12;368(6496):1274–8.

26. Barnes CO, Jette CA, Abernathy ME, Dam KMA, Esswein SR, Gristick HB, et al. SARS-CoV-2 neutralizing antibody structures inform therapeutic strategies. Nature. 2020 Dec;588(7839):682–7.

27. Premkumar L, Segovia-Chumbez B, Jadi R, Martinez DR, Raut R, Markmann AJ, et al. The receptor-binding domain of the viral spike protein is an immunodominant and highly specific target of antibodies in SARS-CoV-2 patients. Sci Immunol. 2020 Jun 26;5(48).

28. Ahmed S, Khan MS, Gayathri S, Singh R, Kumar S, Patel UR, et al. A Stabilized, Monomeric, Receptor Binding Domain Elicits High-Titer Neutralizing Antibodies Against All SARS-CoV-2 Variants of Concern. Front Immunol. 2021;12:765211.

29. Malladi SK, Patel UR, Rajmani RS, Singh R, Pandey S, Kumar S, et al. Immunogenicity and Protective Efficacy of a Highly Thermotolerant, Trimeric SARS-CoV-2 Receptor Binding Domain Derivative. ACS Infect Dis. 2021 Aug 13;7(8):2546–64.

30. Malladi SK, Singh R, Pandey S, Gayathri S, Kanjo K, Ahmed S, et al. Design of a highly thermotolerant, immunogenic SARS-CoV-2 spike fragment. J Biol Chem. 2021;296:100025.

31. Thuluva S, Paradkar V, Gunneri SR, Yerroju V, Mogulla R, Turaga K, et al. Evaluation of safety and immunogenicity of receptor-binding domain-based COVID-19 vaccine (Corbevax) to select the optimum formulation in open-label, multicentre, and randomised phase-1/2 and phase-2 clinical trials. EBioMedicine. 2022 Sep;83:104217.

32. Tai W, Zhang X, Drelich A, Shi J, Hsu JC, Luchsinger L, et al. A novel receptor-binding domain (RBD)-based mRNA vaccine against SARS-CoV-2. Cell Res. 2020 Oct 1;30(10):932–5.

33. Yang J, Wang W, Chen Z, Lu S, Yang F, Bi Z, et al. A vaccine targeting the RBD of the S protein of SARS-CoV-2 induces protective immunity. Nature. 2020 Oct 22;586(7830):572–7.

34. Jungreis I, Sealfon R, Kellis M. SARS-CoV-2 gene content and COVID-19 mutation impact by comparing 44 Sarbecovirus genomes. Nat Commun. 2021 May 11;12(1):2642.

35. Yuan M, Wu NC, Zhu X, Lee CCD, So RTY, Lv H, et al. A highly conserved cryptic epitope in the receptor binding domains of SARS-CoV-2 and SARS-CoV. Science. 2020 May 8;368(6491):630–3.

36. Tortorici MA, Czudnochowski N, Starr TN, Marzi R, Walls AC, Zatta F, et al. Broad sarbecovirus neutralization by a human monoclonal antibody. Nature. 2021 Sep;597(7874):103–8.

37. Liu L, Iketani S, Guo Y, Reddem ER, Casner RG, Nair MS, et al. An antibody class with a common CDRH3 motif broadly neutralizes sarbecoviruses. Sci Transl Med. 2022 May 25;14(646):eabn6859.

38. Yuan M, Zhu X, He WT, Zhou P, Kaku CI, Capozzola T, et al. A broad and potent neutralization epitope in SARS-related coronaviruses. Proc Natl Acad Sci U S A. 2022 Jul 19;119(29):e2205784119.

39. Martinez DR, Schaefer A, Gobeil S, Li D, De la Cruz G, Parks R, et al. A broadly neutralizing antibody protects against SARS-CoV, pre-emergent bat CoVs, and SARS-CoV-2 variants in mice. bioRxiv. 2021 Apr 28;

40. Pinto D, Park YJ, Beltramello M, Walls AC, Tortorici MA, Bianchi S, et al. Cross-neutralization of SARS-CoV-2 by a human monoclonal SARS-CoV antibody. Nature. 2020 Jul;583(7815):290–5.

41. Cohen AA, van Doremalen N, Greaney AJ, Andersen H, Sharma A, Starr TN, et al. Mosaic RBD nanoparticles protect against challenge by diverse sarbecoviruses in animal models. Science (1979). 2022 Aug 5;377(6606).

42. Zhang Y, Sun J, Zheng J, Li S, Rao H, Dai J, et al. Mosaic RBD Nanoparticles Elicit Protective Immunity Against Multiple Human Coronaviruses in Animal Models. Advanced Science. 2024 Mar 17;11(9).

43. Cohen AA, van Doremalen N, Greaney AJ, Andersen H, Sharma A, Starr TN, et al. Mosaic RBD nanoparticles protect against challenge by diverse sarbecoviruses in animal models. Science. 2022 Aug 5;377(6606):eabq0839.

44. Walls AC, Miranda MC, Schäfer A, Pham MN, Greaney A, Arunachalam PS, et al. Elicitation of broadly protective sarbecovirus immunity by receptor-binding domain nanoparticle vaccines. Cell. 2021 Oct 14;184(21):5432–5447.e16.

45. Martinez DR, Schäfer A, Leist SR, De la Cruz G, West A, Atochina-Vasserman EN, et al. Chimeric spike mRNA vaccines protect against Sarbecovirus challenge in mice. Science. 2021 Aug 27;373(6558):991–8.

46. Martinez DR, Schäfer A, Gavitt TD, Mallory ML, Lee E, Catanzaro NJ, et al. Vaccine-mediated protection against Merbecovirus and Sarbecovirus challenge in mice. Cell Rep. 2023 Oct 31;42(10):113248.

47. Zhang Y, Sun J, Zheng J, Li S, Rao H, Dai J, et al. Mosaic RBD Nanoparticles Elicit Protective Immunity Against Multiple Human Coronaviruses in Animal Models. Adv Sci (Weinh). 2024 Mar;11(9):e2303366.

48. Lee DB, Kim H, Jeong JH, Jang US, Jang Y, Roh S, et al. Mosaic RBD nanoparticles induce intergenus cross-reactive antibodies and protect against SARS-CoV-2 challenge. Proc Natl Acad Sci U S A. 2023 Jan 24;120(4):e2208425120.

49. Starr TN, Czudnochowski N, Liu Z, Zatta F, Park YJ, Addetia A, et al. SARS-CoV-2 RBD antibodies that maximize breadth and resistance to escape. Nature. 2021 Sep;597(7874):97–102.

50. Starr TN, Zepeda SK, Walls AC, Greaney AJ, Alkhovsky S, Veesler D, et al. ACE2 binding is an ancestral and evolvable trait of sarbecoviruses. Nature. 2022 Mar;603(7903):913–8.

51. Letko M, Marzi A, Munster V. Functional assessment of cell entry and receptor usage for SARS-CoV-2 and other lineage B betacoronaviruses. Nat Microbiol. 2020 Apr;5(4):562–9.

52. Lee J, Zepeda SK, Park YJ, Taylor AL, Quispe J, Stewart C, et al. Broad receptor tropism and immunogenicity of a clade 3 sarbecovirus. Cell Host Microbe. 2023 Dec 13;31(12):1961–1973.e11.

53. Roelle SM, Shukla N, Pham AT, Bruchez AM, Matreyek KA. Expanded ACE2 dependencies of diverse SARS-like coronavirus receptor binding domains. PLoS Biol. 2022 Jul;20(7):e3001738.

54. Ge XY, Li JL, Yang XL, Chmura AA, Zhu G, Epstein JH, et al. Isolation and characterization of a bat SARS-like coronavirus that uses the ACE2 receptor. Nature. 2013 Nov 28;503(7477):535–8.

55. Liu K, Pan X, Li L, Yu F, Zheng A, Du P, et al. Binding and molecular basis of the bat coronavirus RaTG13 virus to ACE2 in humans and other species. Cell. 2021 Jun 24;184(13):3438–3451.e10.

56. Zhou H, Chen X, Hu T, Li J, Song H, Liu Y, et al. A Novel Bat Coronavirus Closely Related to SARS-CoV-2 Contains Natural Insertions at the S1/S2 Cleavage Site of the Spike Protein. Curr Biol. 2020 Jun 8;30(11):2196–2203.e3.

57. Pramanick I, Sengupta N, Mishra S, Pandey S, Girish N, Das A, et al. Conformational flexibility and structural variability of SARS-CoV2 S protein. Structure. 2021 Aug 5;29(8):834–845.e5.

58. Chakrabarti P, Chakrabarti S. C--H…O hydrogen bond involving proline residues in alpha-helices. J Mol Biol. 1998 Dec 11;284(4):867–73.

59. Prajapati RS, Das M, Sreeramulu S, Sirajuddin M, Srinivasan S, Krishnamurthy V, et al. Thermodynamic effects of proline introduction on protein stability. Proteins. 2007 Feb 1;66(2):480–91.

60. Weber E, Walter E, Pfleiderer T. [Behavior of serum triglycerides, cholesterol, uric acid and sugar levels and of thrombocyte function in heart infarct patients during a 2-year therapy with acetylsalicylic acid, placebo or phenprocoumon]. Verh Dtsch Ges Inn Med. 83:1731–5.

61. van Bergen J, Camps MG, Pardieck IN, Veerkamp D, Leung WY, Leijs AA, et al. Multiantigen pan-sarbecovirus DNA vaccines generate protective T cell immune responses. JCI Insight. 2023 Nov 8;8(21).

62. Kanjo K, Chattopadhyay G, Malladi SK, Singh R, Jayatheertha S, Varadarajan R. Biophysical Correlates of Enhanced Immunogenicity of a Stabilized Variant of the Receptor Binding Domain of SARS-CoV-2. J Phys Chem B. 2023 Mar 2;127(8):1704–14.

63. Khan MS, Jakob V, Singh R, Rajmani RS, Kumar S, Lemoine C, et al. Enhancing Immunogenicity of a Thermostable, Efficacious SARS-CoV-2 Vaccine Formulation through Oligomerization and Adjuvant Choice. Pharmaceutics. 2023 Dec 12;15(12).

64. Mittal N, Kumar S, Rajmani RS, Singh R, Lemoine C, Jakob V, et al. Enhanced protective efficacy of a thermostable RBD-S2 vaccine formulation against SARS-CoV-2 and its variants. NPJ Vaccines. 2023 Oct 25;8(1):161.

65. Roytenberg R, García-Sastre A, Li W. Vaccine-induced immune thrombotic thrombocytopenia: what do we know hitherto? Front Med (Lausanne). 2023;10:1155727.

66. Greinacher A, Langer F, Makris M, Pai M, Pavord S, Tran H, et al. Vaccine-induced immune thrombotic thrombocytopenia (VITT): Update on diagnosis and management considering different resources. J Thromb Haemost. 2022 Jan;20(1):149–56.

67. Sumida SM, Truitt DM, Kishko MG, Arthur JC, Jackson SS, Gorgone DA, et al. Neutralizing antibodies and CD8+ T lymphocytes both contribute to immunity to adenovirus serotype 5 vaccine vectors. J Virol. 2004 Mar;78(6):2666–73.

68. Barouch DH, Pau MG, Custers JHH V, Koudstaal W, Kostense S, Havenga MJE, et al. Immunogenicity of recombinant adenovirus serotype 35 vaccine in the presence of pre-existing anti-Ad5 immunity. J Immunol. 2004 May 15;172(10):6290–7.

69. Wang WC, Sayedahmed EE, Mittal SK. Significance of Preexisting Vector Immunity and Activation of Innate Responses for Adenoviral Vector-Based Therapy. Viruses. 2022 Dec 6;14(12).

70. Fausther-Bovendo H, Kobinger GP. Pre-existing immunity against Ad vectors: humoral, cellular, and innate response, what’s important? Hum Vaccin Immunother. 2014;10(10):2875–84.

71. Chattopadhyay G, Varadarajan R. Facile measurement of protein stability and folding kinetics using a nano differential scanning fluorimeter. Protein Sci. 2019 Jun;28(6):1127–34.

72. Tang J, Zeng C, Cox TM, Li C, Son YM, Cheon IS, et al. Respiratory mucosal immunity against SARS-CoV-2 after mRNA vaccination. Sci Immunol. 2022 Oct 28;7(76):eadd4853.

73. Ye T, Jiao Z, Li X, He Z, Li Y, Yang F, et al. Inhaled SARS-CoV-2 vaccine for single-dose dry powder aerosol immunization. Nature. 2023 Dec;624(7992):630–8.

